# Mapping the bacterial ecology on the phyllosphere of grass hay and the potential hazards of soaking fodder for horse gut health

**DOI:** 10.1101/494799

**Authors:** Meriel JS Moore-Colyer, Annette Longland, Patricia Harris, Susan Crosthwaite

**Affiliations:** Royal Agricultural University, Cirencester Gloucestershire, UK GL7 6JS.; Equine and Livestock Nutrition Services, Pantafallen Fach, Tregaron, Ceredigion, Wales. SY25 6NF; Mars Horsecare UK LTD; Equine Studies Group, Waltham Centre for Pet Nutrition, Waltham-on-the Wolds, Leicestershire, UK. LE14 4RT; Faculty of Biology, Medicine and Health, University of Manchester, UK M13 9PT.

**Keywords:** hay, horse, forage, bacteria, soaking

## Abstract

Globally hay is the preferred forage for stabled horses. Variable nutritional and hygienic quality stimulates pre-feeding soaking to reduce dust and nutrients to reduce respiratory and metabolic disorders in horses. However, this practice has potential negative impacts on horse health. The objectives of this study were to map the bacterial profile of different hays and determine how soaking alters this with the aim of recommending best practice when feeding fodder to stabled horses. Two meadow and one Perennial Ryegrass hays were soaked for 0, 1.5, 9 or 16 hours. Post treatment, hays were analysed for water-soluble carbohydrate (WSC) and total aerobic bacteria (TVC), with differences determined using ANOVA and least significant difference. Bacteria were identified *via* genomic DNA extraction (V3 and V4 variable region of the 16S rRNA gene) and 16S library preparation according to the Illumina protocol. Differences in phyla and family operational taxonomic units within hay types were identified *via* paired t-tests on the DESeq2 normalised data and false discovery rates accounted for using Padj (P<0.05). Mean WSC losses g/kg DM (+/-SE) increased with soaking time being 30 (10.7), 72 (43.7), 80 (38.8) for 1.5, 9 and 16 hours soak respectively. No relationship existed between WSC leaching and bacteria content or profile. Grass type influenced bacterial profiles. Soaking altered the epiphytic bacterial profile across all hays and 9 hours soaking increased richness and Shannon diversity indices. Clustering of bacteria was seen between meadow hays which differed from perennial rye grass and this difference increased post soaking. The normal industry practice of soaking hay for 9 hours pre-feeding cannot be recommended as it increases total bacteria content with noted increases of some potential pathogens. The alterations in bacteria profile and hygienic quality may explain why changing fodder or pre-feeding treatments can frequently precipitate colic in horses.

## Introduction

Grass conserved as hay is an ubiquitous fodder used to feed a wide range of livestock and is still the preferred long forage for stabled equids across the world. [1,2] However, especially in temperate climates, it is difficult to make good quality hay that has low dust, bacteria and mould spore counts. Grass conserved as silage or haylage, which are fermented forages that require less field-drying time [3] are often suggested as suitable alternatives to hay but in many cases these forages are not an economical or practical solution. Small 15-20 kg bales of haylage are expensive and big bales of 200kg or more need mechanical handling and have a shelf-life of 5 days before aerobic despoliation makes it unsuitable to feed. Moreover, the perceived advantages of haylage as being low in dust and lower in non-structural carbohydrate content than hays is not always found. [4]

The nutritional value and hygienic quality of hay depends on a plethora of factors such as grass species, edaphic and environmental conditions during growth and at harvest, maturity at harvest and storage conditions.[4,5]. This makes hay not only a highly variable feed source in terms of nutritional content but can also present hidden challenges to the health of humans handling it and the animals consuming it.

Farmers Lung and Equine Asthma are two well documented conditions arising from the airborne respirable dust (ARD) that is inherent in hay [6]. Veterinary surgeons and horse owners of laminitic or obese horses are also aware that some well conserved hays can contain WSC contents in excess of 310 g/kg DM [7] which although not an issue for horses with high energy demands, makes the hay unsuitable to feed to animals pre-disposed to laminitis who should be fed a forage of less than 100g/kg DM WSC [8]. Both of these conditions have led to the global practice of soaking hay for varying lengths of time before feeding, to reduce the negative impact of ARD and high WSC contents.

While the practice has been previously shown to be effective in both of these endeavours [9,10,11,12,13] soaking hay has some well-documented disadvantages such as nutrient and mineral leaching [9,10,11] production of post-soak liquor that is a biological hazard [14,15] and recorded increases of 1.5 to 5 fold of bacteria in post soaked hay samples [16,17]. To date there is no published information on the resident bacterial profile (s) of hays nor what influence soaking might have on such profiles. Hay is still the most common fodder fed globally to stabled equids so it is vitally important to understand how pre-feeding treatments can influence the hygienic quality of the forage.

Bacteria can form large heterogeneous aggregates, reputed to constitute between 30 and 80% of the total bacteria on plant surfaces. Many of these aggregates also harbour fungi,[18,19,20].which may pose an additional threat to the hygienic quality of the fodder and the health of the animals consuming it. Identifying bacterial profiles of conserved forage will provide new insights as to which bacterial families commonly colonize cut fodder and if these are a potential threat to animal health. Furthermore, a greater understanding of microbial profiles and interactions between bacteria and the phyllosphere of herbage in fodders may help farmers and horse owners decide on the best conservation process and pre-feeding treatments to apply to the fodder.

A series of studies by Dougal *et al.,*[21,22,23,24] has shed light on the bacterial profile, stability and size of the equid gut microbiome. A persistent (at least 6 weeks) core bacterial community was recorded within all regions of the hindgut, but no clustering was seen between individuals (β diversity) according to diet. Diet seemed to influence bacterial profiles within individual horses (α diversity) in that each horse responded differently. Furthermore, the core community has been reported to be smaller than found in ruminants and humans and to be composed of many OTUs of low abundance. Ericsson *et al* [25] found much variation between individuals in foregut bacterial profiles but that hind gut profiles were more uniform. Thus, it would seem that individual horses have a phylogenetic community specific to themselves and their diet and that profile could make them more or less susceptible to dysbiosis when dietary changes are made.

Over 70% of an animal’s immune system is held in the gut-associated lymphoid tissue (GALT) and for an individual to remain healthy and productive their digestive system must also be healthy [26,27]. This is important for horses, as their immune system is continually challenged by frequent mixing with conspecifics at competitions and leisure rides. The economic impact of days lost for training and veterinary bills are obvious, but the positive aspects of long-term animal welfare and public perception of the industry should not be ignored.

The objectives of the current study were to map the resident bacterial profile (s) in different types of hay when dry and to determine how that profile altered post-soaking. Additionally, the study also examined if any relationship existed between WSC leaching and the growth of potentially pathogenic bacteria in post-soaked hay and if such treatment of hay is likely to have a negative impact on the digestive health of the horse. The aim of this research is to reveal unknown information on how pre-feeding treatments affect forage hygiene and thus add to best practice recommendations on feeding fodder to stabled horses.

## Materials and Method

### Hay and sampling procedure

Three replicate bales of three different types of hay, were sourced from 2 farms in Wiltshire, UK. Hays were made in July 2013, were well conserved and had no visible signs of microbial spoilage. Hay types were Perennial ryegrass (*Lolium perenne*) (PRG) and Meadow hay medium cut (MC) (Peploe, Swindon, Wiltshire) and Meadow hay medium cut (MS) (Sian, Great Somerford, Wiltshire). Post-purchase, all bales were labelled and stored off the ground on pallets in a wooden building at the Royal Agricultural University. Each hay type was subjected to the following procedure:

The three replicate bales of each hay type were opened and individually thoroughly with gloved hands on a clean plastic sheet in a glass-house. Two kg of hay was placed into each of 12 small-holed hay nets. Hay nets were then put into purpose made, pre-labelled, polyester hay bags (Haygain Ltd, Hungerford, Berkshire UK) 6 hay nets per bag, and stored until treated. The remaining hay approximately 30 kg was stored in polyester hay bags for later use for bacterial DNA extraction.

### Treatments

Three replicate hay nets for each type of hay were individually subjected to one of the following treatments (3 hays x 4 treatments x 3 replicates n= 36). 1. Dry (D) where no additional treatment was applied to the hay; 2. Soaked (W1.5) by total immersion in 30 litres of clean tap water at 16°C for 1.5 hours, then hung up to drain for 10 minutes; 3. Soaked (W9) as above for 9 hours; 4. Soaked (W16) as above for 16 hours.

Post-treatment, the hay was mixed and two sub-samples were taken. Subsample 1 (approximately 10 g) was placed into a sterile plastic bag and placed in a laminar-flow cabinet (Bassaire, Duncan Rd, Swanwick, Southampton), for bacterial culturing. Sub-sample 2 (approximately 800 g), was weighed onto a pre-weighed foil tray and placed in a forced draught oven at 60°C and dried until a constant weight was obtained for dry matter (DM) determination. The sample was then milled using a 1093 Cyclotec Sample Mill (Foss Sweden) and 50 g of the dried, milled sample was retained and stored in sterile plastic tubes (VWR, UK) for subsequent WSC analyses.

### Bacterial culturing and enumeration

Immediately post-treatment, sub-sample 1 was roughly chopped into 2cm lengths with scissors, (previously wiped with ethanol, and allowed to dry) and thoroughly mixed. A one gram sub-sample was then weighed into a sterile plastic bag (Seward BA6040) to which 79 ml of sterile peptone saline solution (MRD) was added. The bag was then placed into a Lab Blender 80 model (Steward Laboratory, Blackfriars Rd, London). The mixture was then ‘blended’ for 2 minutes in order to wash bacteria from the hay into the solution as for 3M petrifilms (3M Microbiology, 2013). One millilitre of the blended solution was placed into a sterile screw-cap tube (VWR, UK) containing 9 ml MRD. Serial dilutions were prepared to 10^-6^. A 1 ml sample was then taken from 10^-2^, 10^-4^10^-6^ dilutions and separately placed onto pre-labelled 3M Aerobic TVC 20 cm^2^ petrifilm, (3M Microbiology, Carl-Schurz-Straβe 1, Germany). Petrifilms are a sample ready culture medium, containing nutrients, a cold water-soluble gelling agent and a tetrazolium indicator. Three petrifilms were prepared for each sample and incubated for 3 days at 32°C.

Colony numbers were enumerated using an illuminated magnifier. All vital stained colonies were counted. When colony numbers were particularly dense and small and >100 per film, three representative 1 cm squares were counted. The average was determined, and scaled up 20-fold as an estimation of the count per film.

### Water soluble carbohydrate analyses

Immediately post-treatment, approximately 300 g of hay was weighed out into pre-weighed foil trays. These were placed into a forced-draught oven and dried for a minimum of 48 hours at 65°C until constant weight was reached. Post-drying samples were milled through 0.75 mm mesh and re-bagged into 100g DM sub-sample batches. Water soluble carbohydrate (WSC) analyses was then carried out on 3 replicates per sample using the Phenol-sulphuric acid method ^(28)^.

### Preparation of hay for DNA extraction and sequencing

The remaining 30 kg of stored dry hay from each hay type was sub-sampled 3 times, taking approximately 100 g for each sample. Each of the three replicate samples underwent the following procedure. A 0.5g sub-sample was placed in a 50 ml glass tube. Seven and a half ml of tap water at 16 °C was added to each tube, covered with foil and placed in an incubator at 16°C. After soaking, for 0, 1.5, 9 or 16 hours, samples were placed on Whatman filter paper for 10 minutes and then chopped into approximately 0.5 cm lengths for DNA extraction. Genomic DNA was extracted using the MoBio PowerMaxSoil TM kit (MoBio, Carlsbad, CA, USA) according to the manufacturer’s protocol. 16S library preparation was carried out according to Illumina’s protocol. Briefly, genomic DNA was amplified with forward primer 5’-TCGTCGGCAGCGTCAGATGTGTATAAGAGACAGCCTACGGGNGGCWGCAG-3’ and reverse primer 5’ - GTCTCGTGGGCTCGGAGATGTGTATAAGAGACAGGACTACHVGGGTATCTAATCC-3’ targeting the V3 and V4 variable region of the 16S rRNA gene. Twenty five microlitre PCR reactions contained 12.5 μl 5 KAPA HiFi HotStart Ready Mix Master Mix, 5 μM final concentration of forward and reverse primers and 21 ng gDNA. Amplification program: 95 °C 3 mins, 25 cycles of 95 °C 30 s, 55 °C 30 s, 72 °C 30 s, final extension 72 °C 5 min. A subsequent limited-cycle amplification step was performed to add multiplexing indices and Illumina sequencing adapters. Ampure XP beads were used in the PCR clean-up after 1^st^ and 2^nd^ stage PCR. The libraries were then normalised, pooled and sequenced on the MiSeq platform. The quality of raw sequence data was first checked using FastQC. Next, Trimmomatic was used to filter out poor reads, sequences were truncated to 180 bp and paired ends joined using SeqPrep. Sequences were uploaded to QIME where they were clustered into operational taxonomic units with a 97% similarity cut off using as a reference the pre-clustered versions of the Greengenes database. Sample OTUs were merged using a personal Java script and differences tested with DESeq2 [29].

Principal Component Analyses (PCA) and differential counts between conditions were performed. PCA 1 = strongest pattern of variance and PCA 2 the second strongest pattern of variance. Differences between a) the three hay types and b) the four soaking treatments within hay type on the proportion of OTUs of bacterial phyla and family was performed using paired t-tests on the DESeq2 normalised data. In order to take account of false discovery rates that can occur in such data sets with multiple parallel measurements, the Padj value of <0.05 was taken as the cut-off point for significant differences.

### Bacterial diversity, richness and similarity between hay types and treatments

Shannon diversity indices and richness tests were calculated on the *ca* 250 bacterial family OTUs identified. The three un-treated (D) hay types were compared as were the diversity and richness within hays comparing the dry treatment with each of the 1.5, 9 and 16 hours soaking treatments.

Jaccard similarity index was used to determine the level of commonality between the hay types and within individual hays between dry control and soaking treatment. Difference in diversity between the hay types and within hay types compared with soaking time were determined using the Hutchinson’s t-test.

### Data analyses and sample size

Differences in WSC content from this Randomised Block Experiment were determined using analysis of variance (ANOVA), with hay (3), bale (3) and treatment (4) as fixed factors; thus sample size was n = 36. Differences between means were calculated using least significant difference (LSD) test where LSD= t _(error df)_ x s.e.d. Differences in the numbers of bacterial colony forming units (CFU) were determined using ANOVA on log 10 transformed data using Genstat 18 as described by the procedure for right-handed skewed data [30]. Differences between treatment means was determined using least significant difference (LSD) test where LSD= t (error df) x s.e.d. Results for WSC contents were expressed as g/kg on a DM basis, while those for total viable count (TVC) were expressed as geometric mean colony forming units (CFU)/g on an as fed basis, as this value approximates closely to the median [30] which is widely accepted to be the most accurate expression of the distribution of the CFU in the samples.

## Results

### Dry matter, water soluble carbohydrate and microbial content in hay

The three hays, two mixed species meadow hays (MC and MS) and a perennial ryegrass (PRG) hay were grown on 2 different farms in Wiltshire. The MC and MS hays contained similar varieties of grass species inclusive of perennial ryegrass, both rough stalked and annual meadow grass, Timothy, Yorkshire fog, Cocksfoot, and small amounts of Crested Dogs Tail. The MS hay was more mature with a higher proportion of stem to leaf than the MC. The hays were well conserved as all three hays were above the 85% DM recommended to ensure good crop conservation [3] as shown in Table 1.

**Table 1.** WSC content (g/kg DM) and bacterial content (CFU/g) of dry MS, MC and PRG hays.

|  | MC | MS | PRG | sed | Sig |
| --- | --- | --- | --- | --- | --- |
| DM g/kg | 930 | 970 | 930 |  |  |
| WSC (g/kg DM) | 242.6<sup>b</sup> | 124.6<sup>a</sup> | 204.3<sup>b</sup> | 17.76 | 0.006 |
| TVC (log 10 CFU/g) | 8.06 | 6.99 | 7.32 | 0.765 | 0.457 |

<sup>ab</sup> Values in the same row not sharing common superscripts differ significantly (P<0.05)

The WSC contents of the three hays before treatment are detailed in Table 1 and show that MS hay was significantly (P<0.05) lower in WSC content than the other two hays being 118 and 80 g WSC/kg DM lower than MC and PRG respectively.

There was no significant difference in the abundance of bacteria as measured by TVC (CFU/g) between the three dry hays. The geometric mean (Table 2) CFU/ g revealed high contents of bacteria for all the hays according to the classification used by Adams [31] and the 30 ×10^6^ the CFU/g noted by Bucher and Thalmann [32]. The MS hay had the lowest content of bacteria at 2.4×10^7^ CFU/g and the MC the highest at 7.6×10^8^ CFU/g.

**Table 2.** The effect of 4 different soaking times 0, 1.5, 9 and 16 hours on the dry matter (DM), bacterial content (CFU/g) and water soluble carbohydrate (WSC) in 2 types of mixed species meadow hay (MS and MC) and perennial rye grass (PRG) hay for horses.

| param<br>eter | MS 0 | MS 1.5 | MS 9 | MS 16 | MC 0 | MC 1.5 | MC 9 | MC 16 | PRG 0 | PRG 1.5 | PRG 9 | PRG 16 |
| --- | --- | --- | --- | --- | --- | --- | --- | --- | --- | --- | --- | --- |
| DM | 97 | 49 | 44 | 46 | 93 | 49 | 47 | 43 | 93 | 37 | 35 | 43 |
| SD | 1.4 | 6 | 3.4 | 1.4 | 1.2 | 2.9 | 4.4 | 3.4 | 1.6 | 5.9 | 3.5 | 12.6 |
| % H<sub>2</sub>O | 3 | 51 | 56 | 54 | 7 | 51 | 50 | 54 | 7 | 60 | 62 | 54 |
| Log<br>TVC | 7.38 | 7.37 | 7.47 | 7.50 | 8.88 | 7.49 | 7.63 | 7.57 | 7.6 | 7.42 | 8.85 | 7.56 |
| SD | 0.78 | 0.15 | 0.05 | 0.18 | 1.15 | 0.09 | 0.08 | 0.23 | 0.57 | 0.0 | 1.01 | 0.03 |
| Geo<br>mean<br>TVC | 24016800 | 23663478 | 29812800 | 31529733 | 762659233 | 31082066 | 42603778 | 36928044 | 37680217 | 26159366 | 715965200 | 35903500 |
| % diff |  | 0.98 | 1.24 | 1.32 |  | 0.04 | 0.06 | 0.05 |  | 0.7 | 19.0 | 0.95 |
| WSC | 125 | 104 | 103 | 97 | 242 | 200 | 148 | 151 | 204 | 166 | 103 | 83 |
| SD | 23.6 | 6.3 | 6.3 | 11.4 | 6.8 | 124 | 13.6 | 13.8 | 21.9 | 20.3 | 3.3 | 4.5 |
| % loss |  | 17 | 18 | 23 |  | 17 | 39 | 38 |  | 19 | 50 | 60 |

### The effect of soaking time on the dry matter, water soluble carbohydrate and microbial content in hay

Post-soaking, the forages absorbed between 50 and 62% additional moisture with no pattern emerging according to soaking time.

All hays lost progressively more WSC up to 9 hours soaking. Table 3 details the average WSC loss across all three hay types to be 33.7 g/kg DM post the 1.5 hours soak (P<0.05); a further drop (P<0.05) of 38 g/kg DM was noted when soaking was increased by 7.5 hours but no further loses were recorded when soaking was extended by a further 7 hours.

**Table 3.**
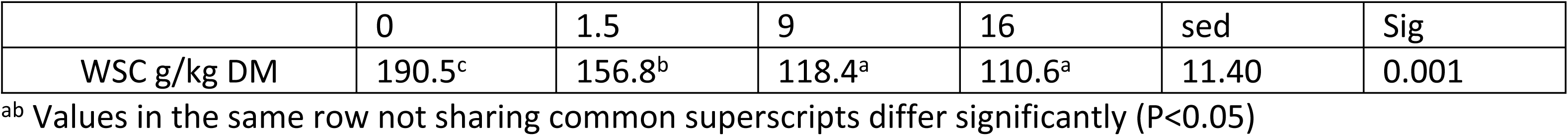
Mean WSC content across all 3 hays after soaking for 0, 1.5, 9 and 16 hours in water at 16°C.

When looking at individual hays and treatments (Table 2) the PRG showed greatest % WSC losses of 19, 50 and 60% for 1.5, 9 and 16 hours soaks respectively, whereas the MS showed the least losses of WSC of 17, 18 and 23%. The hay with the highest starting WSC content showed intermediate losses of 17, 39 and 38% demonstrating that in this study no relationship existed in these three hays between WSC content and WSC leaching across a range of soaking times.

Table 2 Shows that soaking produced a highly variable response in CFU/g across soaking times and hays. Soaking for 9 hours produced a wide range of increases in % of CFU/g in MS and PRG of 1.24 (MS) to 19 (PRG) times that found in the dry samples, whereas a reduction of 6% was recorded for MC hay. Reductions in CFU/g were noted across all hays for the shortest soaking time of 1.5 hours ranging from a 2% in MS to 30% in PRG, but response to 9 and 16 hours soaking were less consistent with some increases at longer soaking times. Therefore, as with WSC levels, the quantitative response of bacteria to soaking in different hays as determined by CFU / g of hay was highly variable and showed no pattern according to soaking time.

### Profile of bacteria in dry hays using 16S rRNA sequencing

Across all three hays a total of 27 phyla and 265 families were identified. All 27 phyla were present in each of the dry hays, although the family proportional profiles differed between the hays. PCA Figure 1 shows degree of similarity between the two meadow hays but the PRG was clearly different. The profile of bacterial phyla are shown in Figure 2 representing proportions of operational taxonomic units (OTUs) found in the three hays when dry. The 4 phyla that represented >0.96 of the bacteria present were *Proteobacteria, Cyanobacteria, Actinobacteria* and *Bacteroidetes* and there were no significant differences between any of the hays for these phyla. The other 23 phyla identified comprised less than 4% of the proportion of OTUs in each hay but differences were noted between MC and MS in the proportions of *Verrucomicrobia* and A*cidobacteria* and between MC and PRG in *Fusobacteria* and *Nitrospirae.*

**Figure 1.**
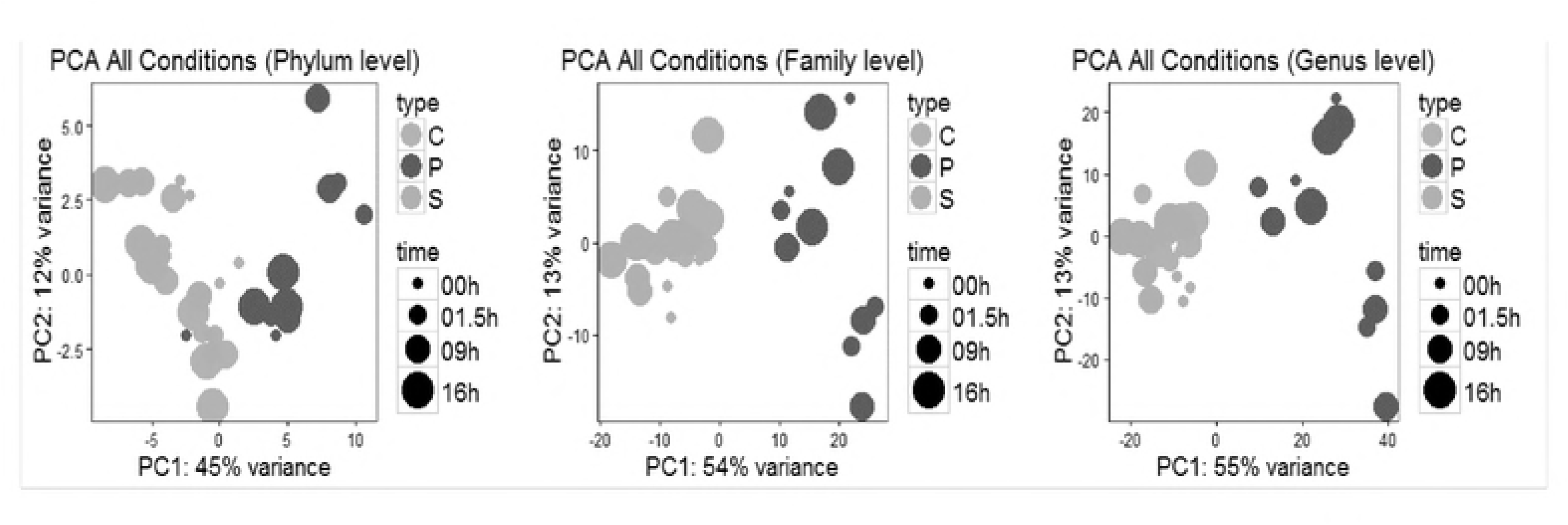
Principal component analyses (PCA) of bacteria identified in Meadow Charlie (MC), Meadow Sian (MS) and Perennial Ryegrass hays when dry (0) soaked for 1.5 hrs, 9 hrs and 16 hrs in water

**Figure 2.**
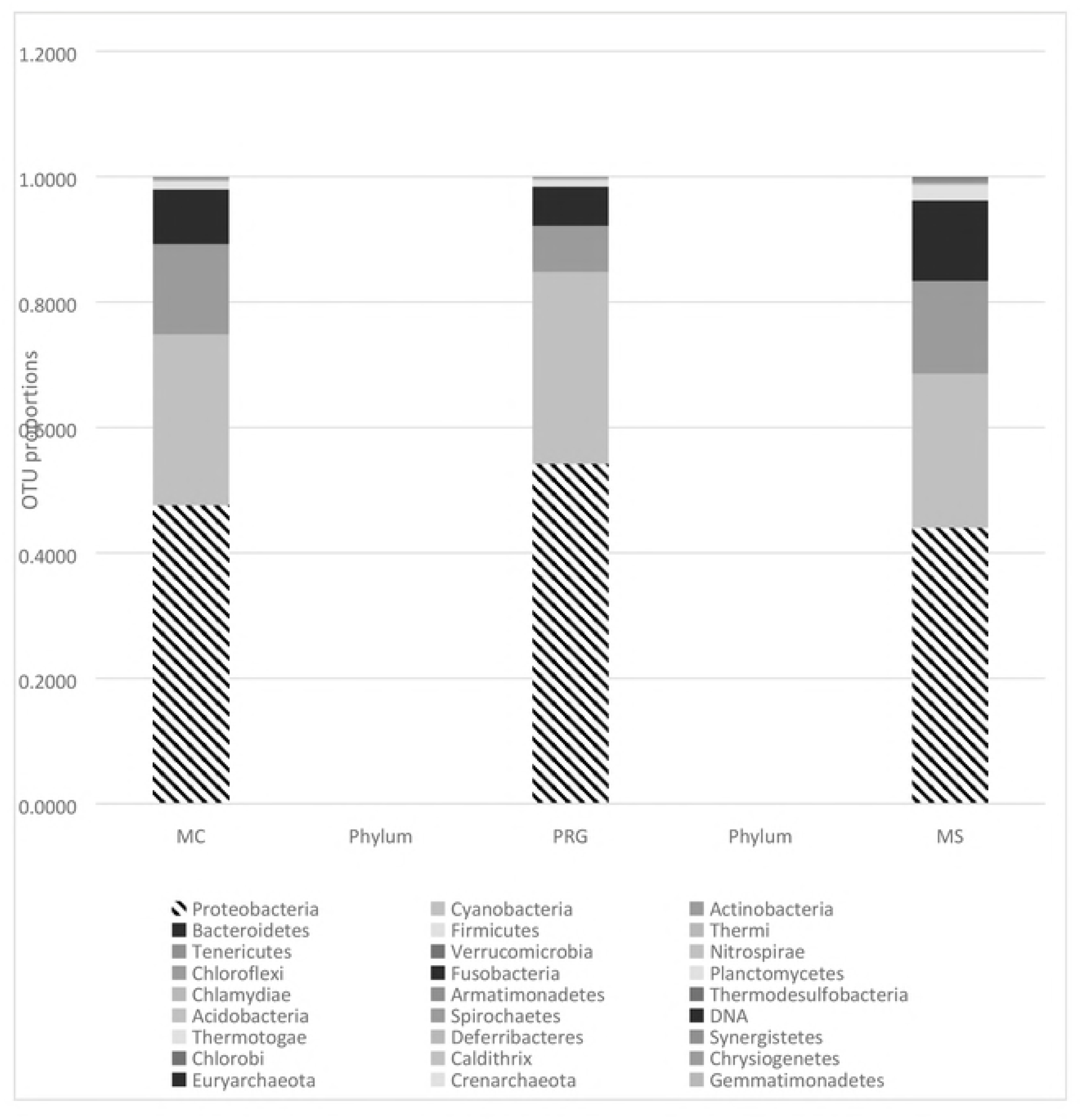
Proportions of operational taxonomic units (OTUs) of bacteria phyla present in dry samples of Meadow Charlie (MC), Meadow Sian (MS) and Perennial Ryegrass (PRG) hays

The profile of OTUs of bacterial families shown in Figure 3 indicates that *Rivulariaceae,* (phylum *Cyanobacteria*) *Sphingomonadaceae Pseudomonadaceae* and *Enterobacteriacea* (all in the phylum *Proteobacteria*) comprised between 0.48 and 0.69 of the bacteria present in all three hays. *Enterobacteriacea* was the only major family present to be higher (P<0.05) at 25% in PRG than in the other two hays, with MC at 0.02 and MS at 0.05 respectively. Differences (P<0.05) between bacterial families that comprised between 31 and 52% of the total present are detailed in Table 4. The two meadow hays were similar with only 3 differences whereas the PRG differed from MC in 12 families and with MS in 20 families.

**Table 4.** Differences (P<0.05) between proportions of bacteria families present in dry Meadow Charlie (MC), Meadow Sian (MS) and Perennial Ryegrass (PRG) hays before treatment.

| MC vs PRG | MS vs MC | PRG vs MS |
| --- | --- | --- |
| Enterobacteriaceae |  | Enterobacteriaceae |
| Xanthomonadaceae |  | Xanthomonadaceae |
| Pasteurellaceae |  | Williamsiaceae |
| Methylocystaceae | Methylocystaceae | Pasteurellaceae |
| Patulibacteraceae |  | Patulibacteraceae |
| Amoebophilaceae | Moraxellaceae | Micrococcaceae |
| Bartonellaceae | Chthoniobacteraceae | Methylophilaceae |
| Micrococcaceae |  | Chromatiaceae |
| Sanguibacteraceae |  | Moraxellaceae |
| Aeromonadaceae |  | Coxiellaceae |
| Francisellaceae |  | Bacteroidaceae |
| Carnobacteriaceae |  | Dermabacteraceae |
|  |  | Sanguibacteraceae |
|  |  | Acidobacteriaceae |
|  |  | Aeromonadaceae |
|  |  | Chthoniobacteraceae |
|  |  | Piscirickettsiaceae |
|  |  | Vibrionaceae |
|  |  | Francisellaceae |
|  |  | Sporichthyaceae |

**Figure 3.**
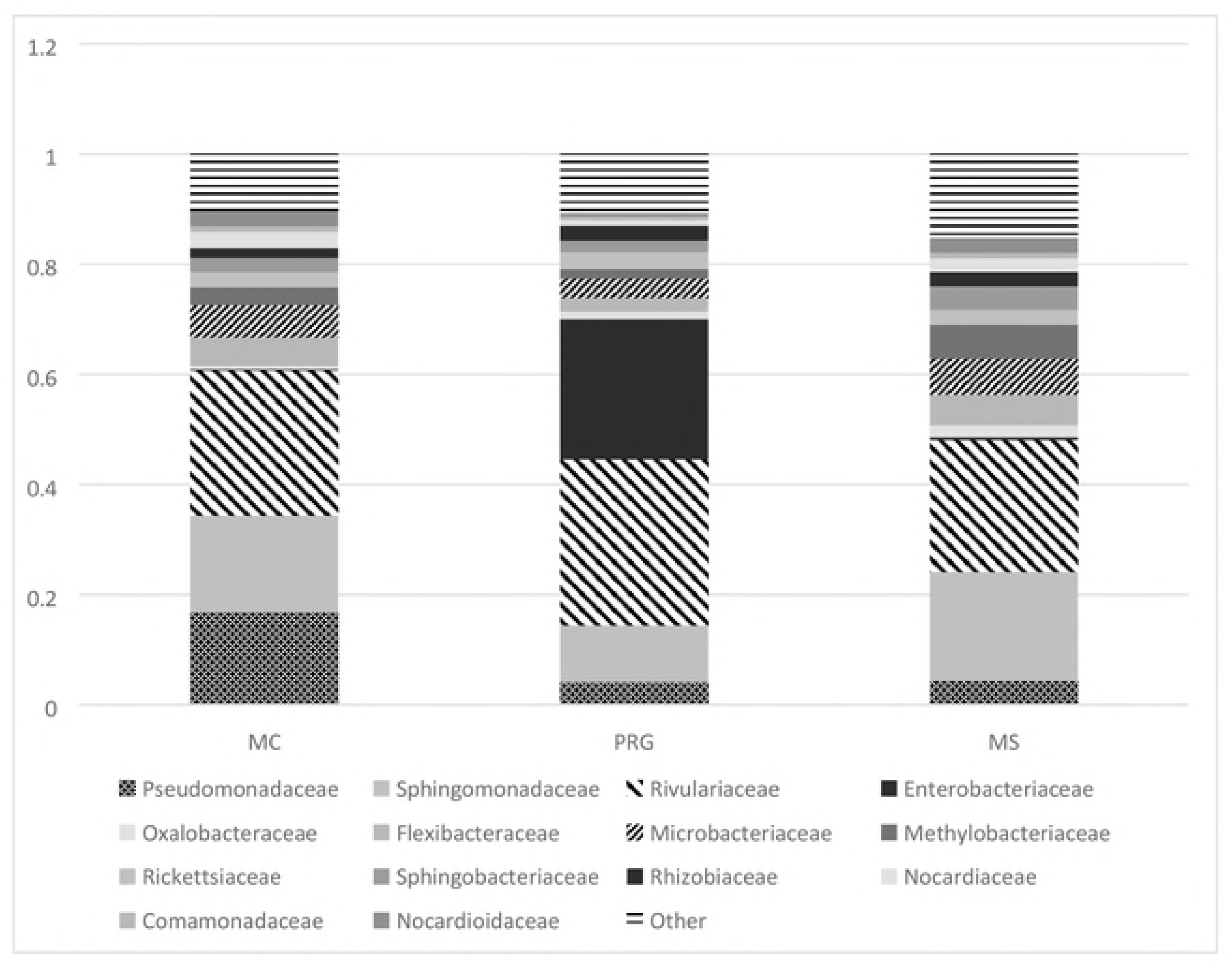
Proportions of operational taxonomic units (OTUs) of bacteria families present in dry samples of Meadow Charlie (MC), Meadow Sian (MS) and Perennial Ryegrass (PRG) hays

### Bacterial family diversity, richness and similarity in 3 dry hays

As detailed in Table 5, the MC hay had the greatest family richness of 230 and a Shannon Diversity Index (H) 2.6; MS was slightly lower at 228 and had a higher H value of 2.8, whereas PRG was lowest at 218 and an H value of 2. The Jaccard Similarity Index (J), shown in Table 6 showed PRG and MC shared 81% of bacterial families, PRG and MS shared 87% and MS and MC shared 86% of the bacterial families sequenced.

**Table 5.** Richness of bacterial families and Shannon Diversity Index (H) within the 3 hays Perennial Ryegrass (PRG), Meadow Charlie (MC) and Meadow Sian (MS) when soaked in water for 0, 1.5, 9 and 16 hours.

| Hay | Richness | H Index |
| --- | --- | --- |
| PRG D | 218 | 2 |
| PRG 1.5 | 215 | 2 |
| PRG 9 | 224 | 3 |
| PRG 16 | 238 | 2.1 |
| MC D | 230 | 2.6 |
| MC 1.5 | 215 | 2.5 |
| MC 9 | 229 | 2.8 |
| MC 16 | 216 | 2.2 |
| MS D | 228 | 2.8 |
| MS 1.5 | 236 | 2.9 |
| MS 9 | 245 | 3 |
| MS 16 | 236 | 2.8 |

**Table 6.** The effect of soaking treatment on the similarity of bacterial families in Perennial Ryegrass (PRG), Meadow Charlie (MC) and Meadow Sian (MS) hays as calculated using the Jaccard Similarity Index.

| Hay comparison | Jaccard Similarity Index |
| --- | --- |
| PRGD vs PRG 1.5 | 81 |
| PRG D vs PRG 9 | 82 |
| PRG D vs PRG 16 | 85 |
| MC D vs MC 1.5 | 87 |
| MC D vs MC 9 | 93 |
| MC D vs MC 16 | 91 |
| MS D vs MS 1.5 | 92 |
| MS D vs MS 9 | 90 |
| MS D vs MS 16 | 91 |

### Effect of soaking for 1.5, 9 and 16 hrs on bacteria phyla and families within hay types

Post soaking (Table 7), the phyla *Armatimonadetes, Cyanobacteria* and *Thermi* all decreased significantly (P<0.05) across all three hays when comparing dry hay with 1.5, 9 and 16 hours soaking, while *Fusobacteria and Acidobacteria* increased across all hay types with soaking. The PRG hay showed more alterations in bacterial phyla as a result of soaking than either of the MC or MS hays.

**Table 7.** Within hay differences in OTU phyla between hay and hay soaked for 1.5, 9 or 16 hours.

|  | MC | PRG | MS |
| --- | --- | --- | --- |
| 0 vs 1.5 | NS | Armatimonadetes | NS |
|  |  | Fusobacteria |  |
|  |  | thermi |  |
| 0 vs 9 | Cyanobacteria | Fusobacteria | Acidobacteria |
|  |  | Thermi | Cyanobacteria |
| 0 vs 16 | NS | NS | Cyanobacteria |
|  |  |  | Fusobacteria |

The effect of soaking on the richness of bacterial families and the H index can be seen in Table 5. Soaking had a variable effect on the richness and H index. In PRG, richness tended to increase with soaking time whereas in both meadow hays the richness at the 9-hour soak was highest with 1.5 and 16 hours being similar. The greatest diversity was noted for MS after the 9 hour soak which gave an H index of 3 and a richness of bacterial families of 245.

Figures 4, 5 and 6 show the alterations in proportions of bacterial families after soaking for 1.5, 9 and 16 hours for MC, PRG and MS hays respectively. Of the four main bacterial families that were present in the dry hay, *Rivulariaceae*, (grey) (phylum *Cyanobacteria*) *Sphingomonadaceae* (orange) *Pseudomonadaceae* (blue) and *Enterobacteriacea* (yellow) (all in the phylum *Proteobacteria*) behaved differently in each of the hays across the different soaking times. While MS and MC hays did show alterations in bacteria post-soaking, the PRG showed greater fluctuations and thus responded more to soaking than the meadow hays.

**Figure 4.**
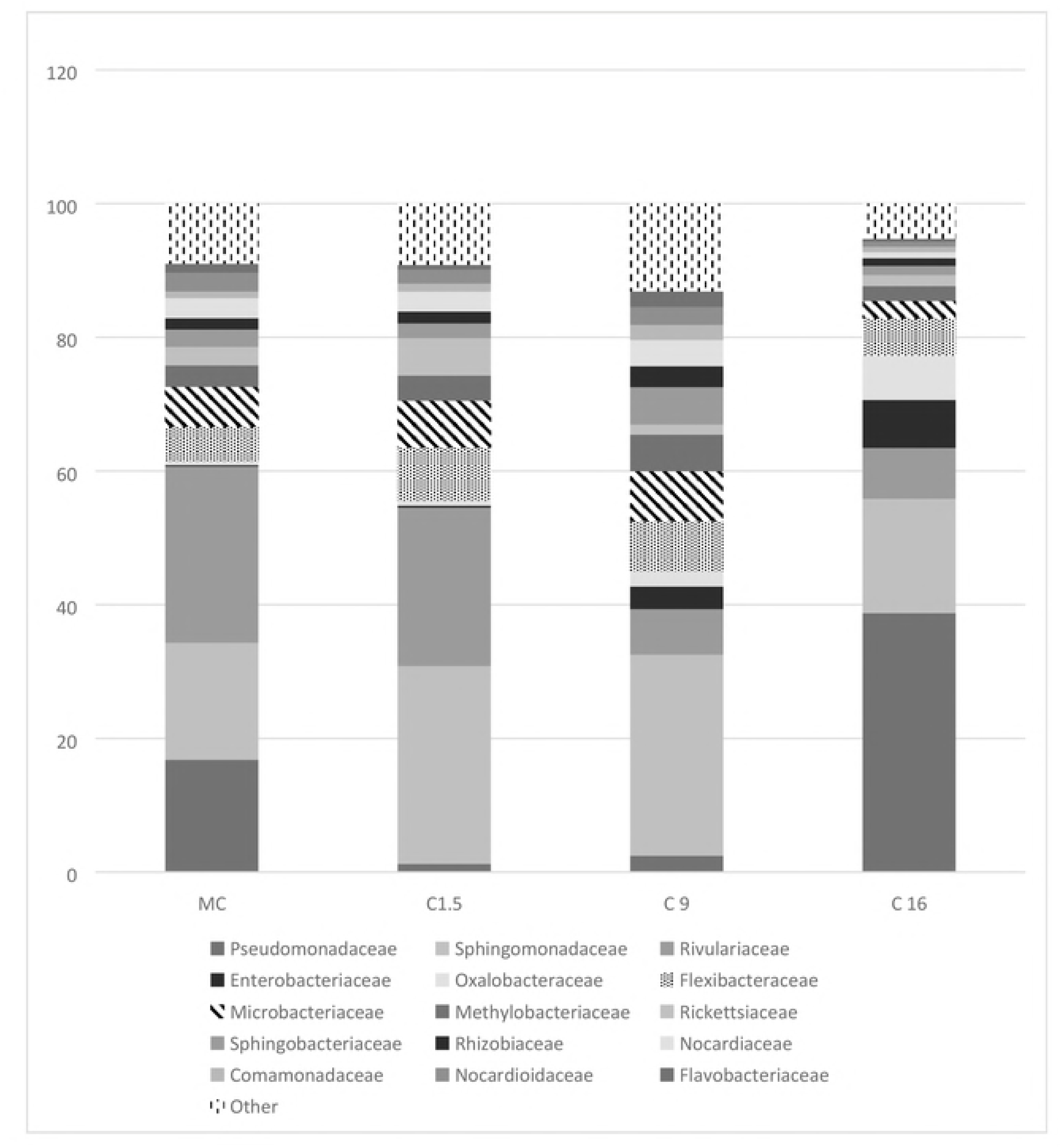
Proportions of operational taxonomic units (OTUs) of bacteria families present in Meadow hay Charlie (MC) when dry, and post soaking in water for 1.5, 9 and 16 hours

**Figure 5.**
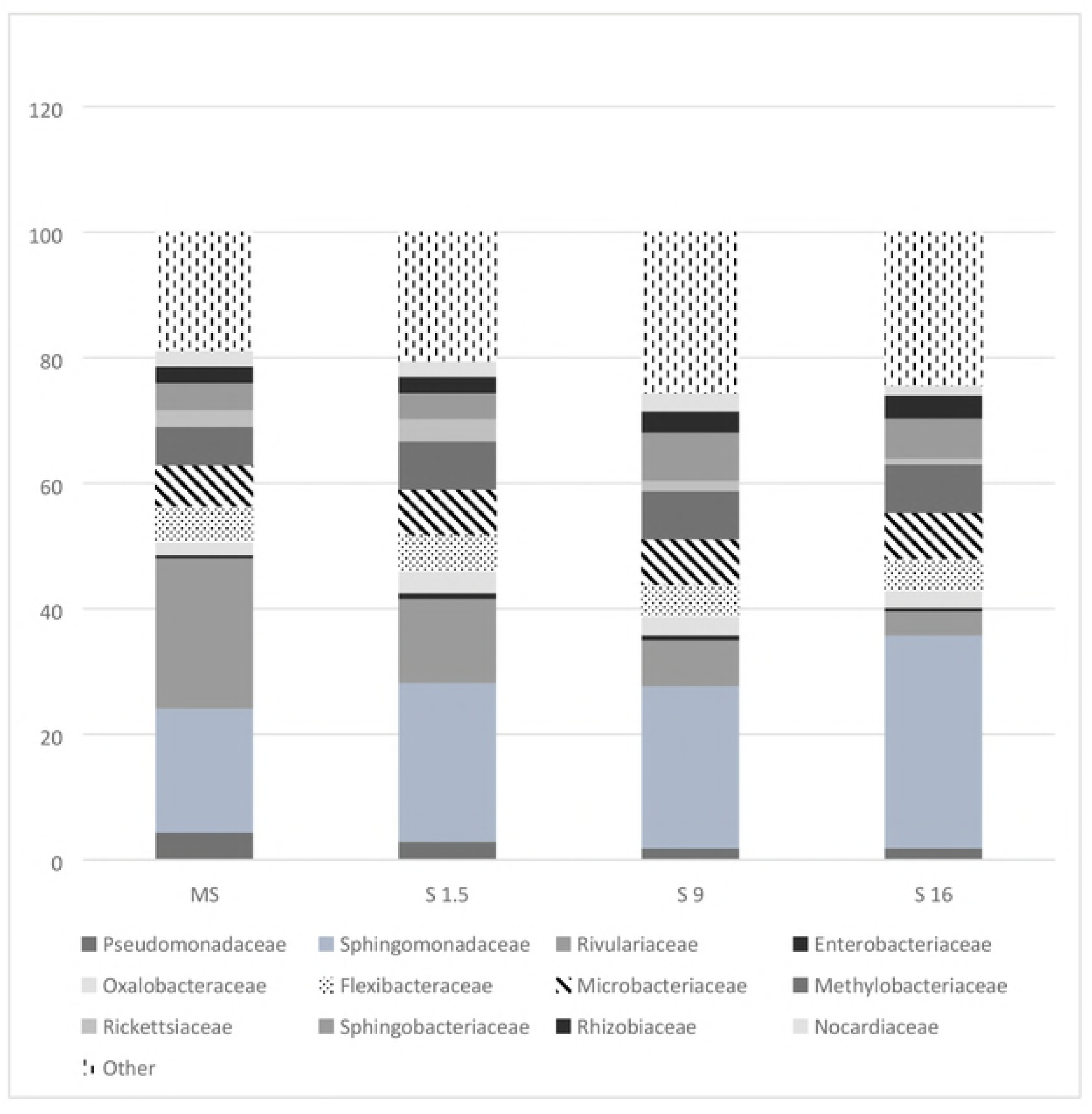
Proportions of operational taxonomic units (OTUs) of bacteria families present in Meadow hay Sian (MS) when dry, and post soaking in water for 1.5, 9 and 16 hours

**Figure 6.**
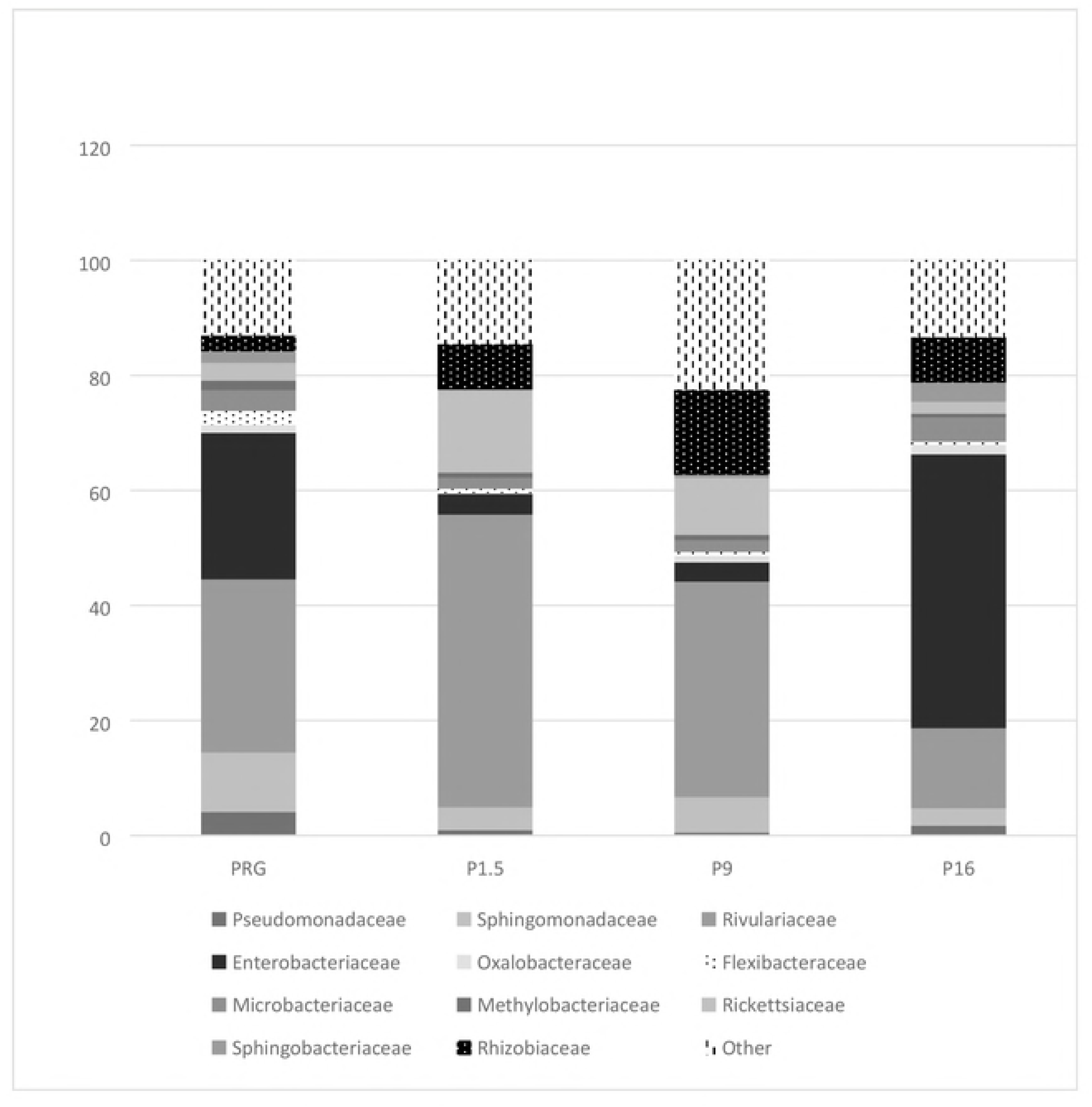
Proportions of operational taxonomic units (OTUs) of bacteria families present in Perennial Ryegrass hay (PRG) when dry, and post soaking in water for 1.5, 9 and 16 hours

*Pseudomonadaceae* proportions decreased in PRG when soaked for 9 hours, but in the other two hay samples for all soaking times no differences were detected for this family. *Sphingomonadaceae* decreased in PRG after1.5 hours of soaking but showed no other variation from the dry hay. Proportions of *Enterobacteriacea* increased in MC hay after 9 and 16 hours soaking but decreased in PRG after soaking for 1.5 and 9 hours but had increased again after 16 hours soaking. *Rivulariaceae,* decreased in both MC and MS when soaked for 9 and 16 hours.

Of the remaining bacterial families that comprised between 31 to 51% of the bacteria present *Xanthomonadaceae*, a family containing important animal and plant pathogens, increased in all hays at 9 hours. Table 8 details the bacterial families that were influenced either positively or negatively by soaking. Those that increased included bacteria that favour aquatic habitats or are fermentative in nature utilising sugar to produce ethanol (*Rhodobacteraceae, Aeromonadaceae Rhodocyclaceae, Gemellaceae, Acetobacteraceae*). Potential pathogens such as *Mycobacteriaceae, Burkholderiaceae, Bacillaceae, Anaplasmataceae, Veillonellaceae, Leptotrichiaceae, Fusobacterium* (a strong biofilm anchor), all increased post soaking but were present in very small proportions they are unlikely to be of clinical importance to the horse.

**Table 8.**
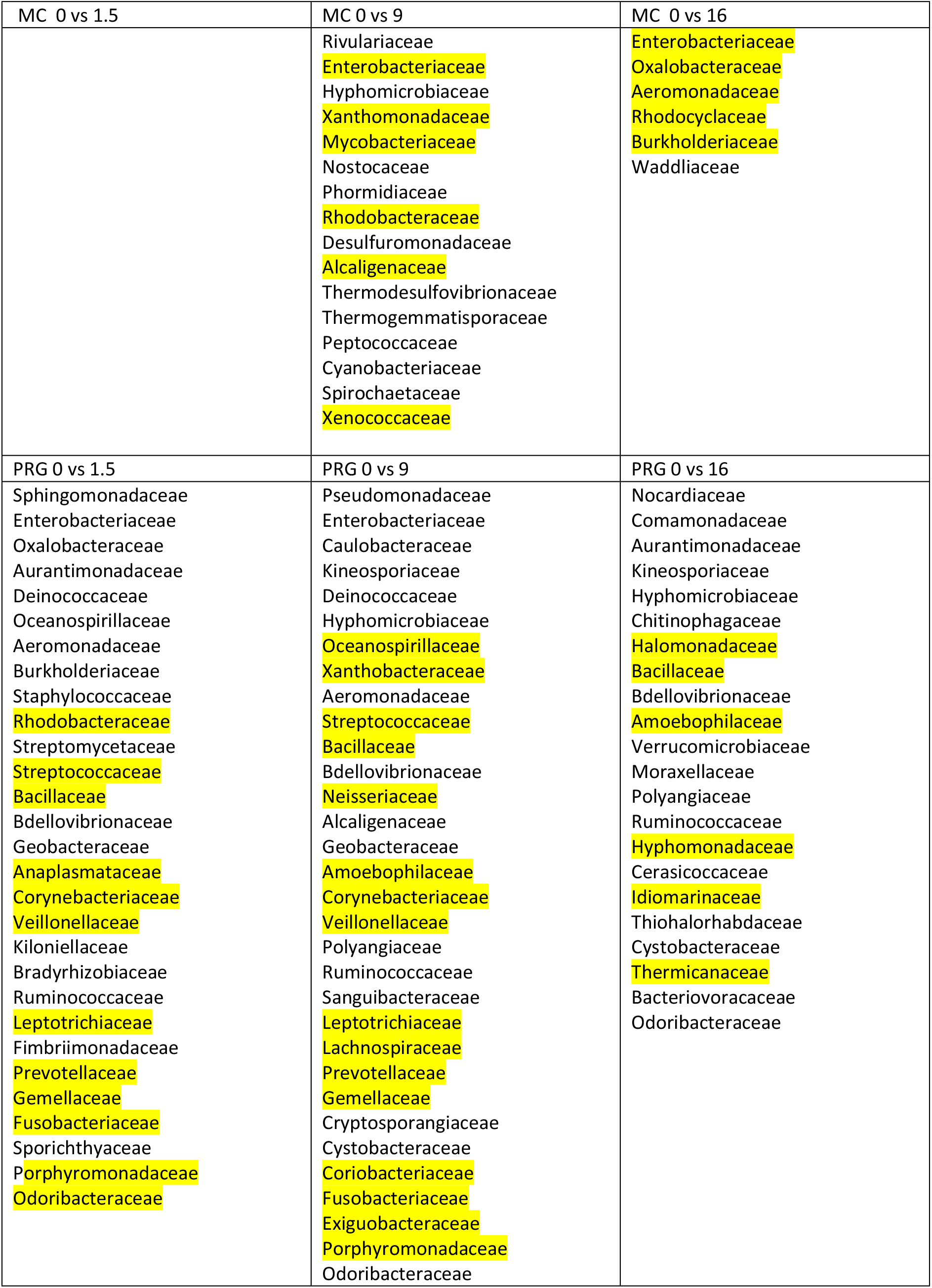
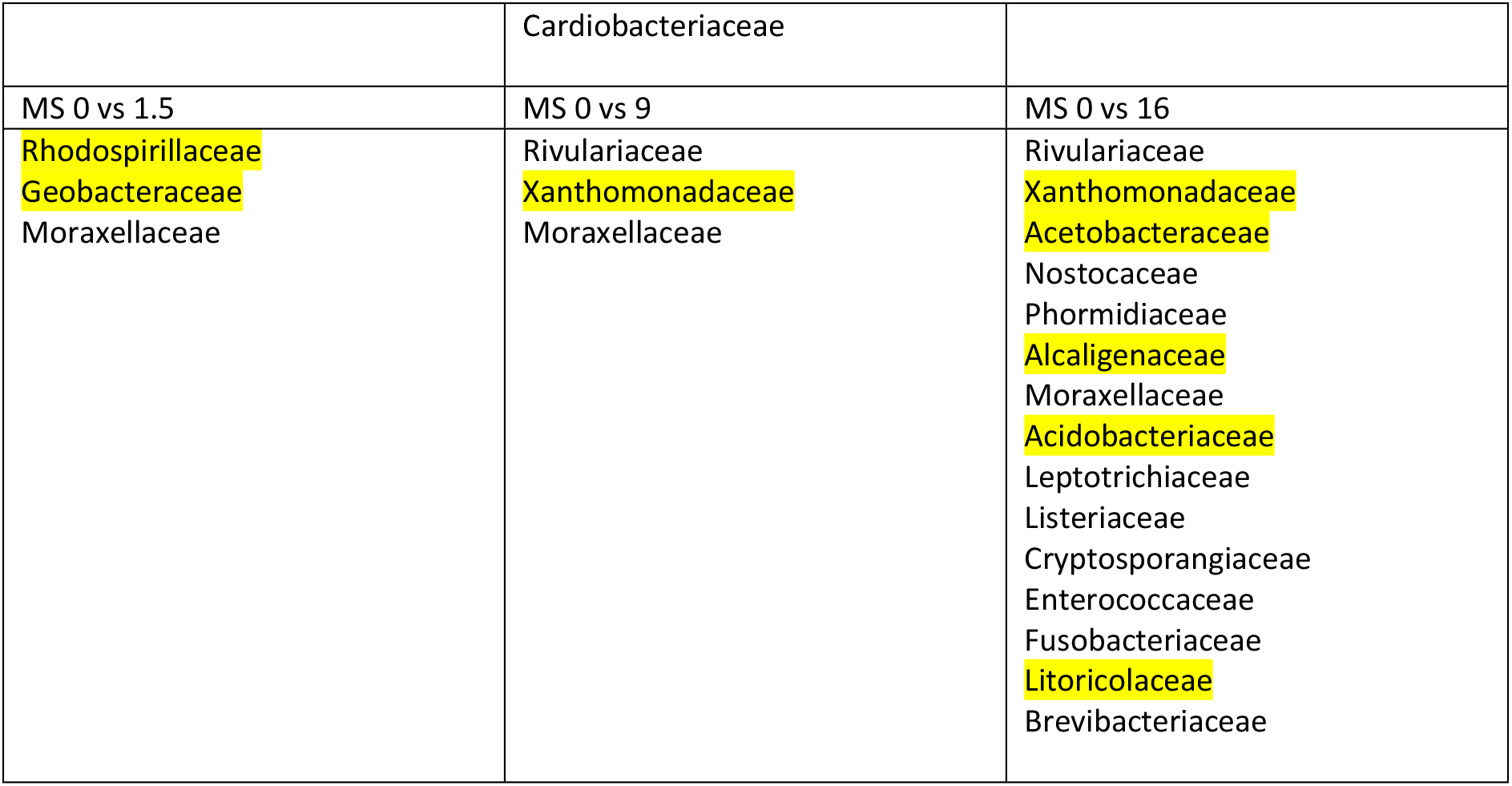
Comparison within each hay type Perennial Ryegrass (PRG), Meadow Charlie (MC) and Meadow Sian (MS) between dry hay and hay soaked for 1.5, 9 and 16 hours within on the increase (highlighted in yellow) or decrease of bacteria families.

## Discussion

### Dry hay, water soluble carbohydrate content and microbial colony forming units (CFU) / g

The range of WSC in the three hays of 125 to 242 g/kg DM is typical of UK hay and agrees with previously published values [11,33]. MC and PRG hays contained 100 to 140g/kg DM more WSC than is currently recommended for forages intended to be fed to equids with a pre-disposition to laminitis and such levels stimulate horse owners to reduce the level of WSC by soaking for extended periods.

Lindow and Brandl [34] noted that bacteria are by far the most numerous colonists of plant leaves, often being found in numbers up to 10^8^ cells/g of leaf [35,36,37]. Although there were no visible signs of aerobic spoilage in any of the hays in this study, the bacterial CFU / g were notably higher than in previously published findings for a range of single and mixed species hays [11,28,39]. It has been noted by Behrendt *et al.* [40] Muller *et al.*[41] that late harvesting can increase the microbial load in forages. Due to poor early season weather conditions all the hays used in this study were not harvested until late July and this may partially explain the high bacterial levels.

### Profile of bacteria in dry hays using 16S rRNA sequencing

Bacteria are by far the most abundant inhabitants of the phyllosphere of plants. While yeasts are active and effective colonizers, filamentous fungi in the form of spores are more transient occupants [1]. Commonly the study of bacteria on the leaves of plants has been driven by their deleterious effect on plant productivity and has been largely restricted to aerobic culturable gram negative bacteria, particularly *Pseudomonas spp. (syringae)* and *Enterobacteriaceae (Erwinia, Pantoea*), which are two of the most ubiquitous bacterial colonizers of the phyllosphere [34]. Despite the importance of bacteria to compromised plant productivity, limited information is available on the bacterial profile of dry fodders or the effect that any pre-feeding treatments might have on that profile. In a study of bacteria on the phyllosphere of grasses growing in extensive grassland, Behrendt *et al.,* [40] found the most prominent 5 genera (phylum in brackets) were *Pseudomonas (Proteobacteria), Stenotrophomonas (Proteobacteria), Pantoea (Proteobacteria) Clavibacter (Actinobacteria) and Curtobacterium(Firmicutes)* Thus, the bacterial families identified from the grass hays in this study and the genera from Behrendt *et al* [40] shared the phylum *Proteobacteria,* and to a minor extent the phyla *Actinobacteria* and *Firmicutes*, with *Cyanobacteria* being a major constituent in PRG hay but not noted as a major presence on the growing grasses. These differences are not surprising as epiphytic bacterial populations can differ in size between plant species and within plants of the same species. Furthermore, changes in bacterial populations can be rapid and are influenced by a wide range of factors such as physiological age, macro and micro environmental conditions and on-leaf microbial interactions [42,43]. Such factors could readily explain the differences between conserved and growing grasses and within species and between forage types noted here. Variations between phyla on the three hays were only seen for the smaller proportional phyla *Verrucomicrobia, Acidobacteria, Fusobacteria and Nitrospirae*. The data on richness and diversity shows the monoculture of PRG supported a less diverse bacterial community than the two meadow hays which had a high degree of similarity. While physical and nutritional conditions accessible to bacteria can account for considerable variations in plant microbial carrying capacity, individual leaf characteristics can also have an effect. PRG has a shiny under leaf and bacterial establishment and maintenance on the phyllosphere can be affected by glossy mutants with the fewest crystalline waxes. Such leaves prove a less effective host for epiphytic bacteria than those with less shiny cuticles [44]. Such results hint at small-scale interactions between the plant and bacterium that are not yet understood.

### Soaking effects

#### The effect of soaking time on the dry matter (DM) and water soluble carbohydrate (WSC) contents of the hays and their bacterial numbers and profiles in hays

The absorption of water of between 50 and 62% noted here were slightly lower than the 73% recorded by Moore-Colyer *et al.,* [11] for a range of Meadow, Timothy and Italian Rye grass hays also soaked for 9 hours.

Extended soaking periods of between 9 and 12 hours have been recommended by Longland, *et al.,* [45] and Muller *et al.,* [46] as a method by which to reduce the WSC content of fodder intended to be fed to horses with insulin resistance, metabolic syndrome, laminitis or obesity. Muller [46] recorded an average WSC loss of 43% after a 12-hour soak and the 18, 38 and 42% losses recorded in this experiment after 1.5, 9 and 16 hours soaking are in agreement with these values. However, such losses cannot be predicted nor relied upon as variability of loss across the hays were 17% to 60% and echo the caution expressed by Longland *et al.,* [45] who recorded variations in WSC leaching from a variety of hays of 9 to 54% after a 16-hour soak. This study reported no additional benefit in terms of WSC leaching when hay was soaked for longer than 9 hours. No pattern was evident between initial WSC content and post-soaking losses in any of the studies and so losses of WSC due to soaking cannot be predicted according to hay species or WSC content. It is important therefore, to highlight to horse owners that WSC losses from soaking hay cannot be set nor predicted according to either soaking time or hay species and so individual hay response must be tested to achieve the optimum soaking time for each hay type.

There was also a highly variable response in bacterial growth (CFU/g) and in phyla and family profile across soaking times and hays. The similarity noted between the meadows hays in the dry samples continued post soaking with MC and MS producing more similar profiles after treatment compared with PRG. Neither the quantity nor diversity of bacterial growth was correlated with WSC content across the hays. A positive correlation between WSC loss and bacterial CFU/g was seen in MS hay, but this was not repeated in either of the other two hays. This may be partly due to the availability of nutrients on the phyllosphere of the different grasses both before soaking and the amount of WSC leached into the water during soaking. Several studies have revealed [47,9,14] that varying amounts of nutrients can be washed from leaves but what influences the degree of leaching is yet to be determined. As small amounts of sugar, about 0.2 to 10µg can support the growth of 10^7^ to 10^8^ bacterial cells/leaf in the growing plant [34] it is easy to see how more readily available sugar from the leaching of 28 to 121 g WSC/kg hay noted here could support considerable bacterial growth during soaking.

Within the complex multifactorial relationship that exists between bacteria and the phyllosphere, there are bacteria that can increase the wettability of leaves by producing compounds with surfactant properties [38]. Fifty percent of the genus of metabolically diverse and wide niche colonizers *Pseudomonas* have been reported [48] to have this ability. One possible explanation for the degree of WSC leaching is an increase in the wettability which allows solubilisation and diffusion of substrates into the water, increasing availability to colonizing bacteria. The *Pseudomoadaceae* family were present in all the hays in this study and their activity could have influenced the phyllosphere making water penetration more effective and thus facilitating the leaching of nutrients during soaking

As the 16S rRNA sequencing is a qualitative identification of bacteria and not a quantitative measurement of CFU/g, the alterations in certain families may have little impact on the nutrient and hygienic quality of the forage. However, of the major families that accounted for a significant proportion of the bacteria, the *Enterobacteriaceae* comprised 25% of the proportion of bacterial families present in PRG. Muller *et al.,*[38] also recorded higher levels 4.9 log ^10^ CFU/ g of *Enterobacteria* in dry hay samples, compared with the same crop conserved as haylage or silage, thus clearly hay supports the growth of this bacterium. The longer soaking times of 9 and 16 hours caused a proportional increase in this family across all the hays and while this is unlikely to have an impact in hays MC and MS the increase in PRG from 25% up to 47% after 16 hours soaking would have a notable impact on the microbial profile of the fodder. Muller *et al.,*[38] also reported a post-treatment increase in *Enterobacteria* in silage, haylage and hay when soaking for 24 hours, but went on to note that the *Enterobacteriaceae* in silage and haylage are generally considered non-pathogenic. However, the family does contain potentially pathogenic species that produce endotoxins which may be associated with diarrhoea in horses [49].

### The effect of soaking hay on the digestive health of horses

Scouring in stabled horses after a change in fodder is a frequent anecdotally reported occurrence. While this may be attributed in part to an alteration in nutrient profile of the feed, poor forage hygiene derived from bacterial and mould proliferation has been associated with colic in horses [50]. A similar effect has been noticed in humans where the presence of pre-harvest epiphytic bacteria on fruit and vegetables has been associated with multiple outbreaks of food-borne illness [51]. Clearly, for both species ingested bacteria survive the low pH of the stomach and are therefore able to colonize and upset the normal microbiome causing dysbiosis.

Although highly variable between horses, Ericsson *et al.*[25] reported an abundance of α-*Proteobacteria* in the upper gastrointestinal tract of 9 healthy horses. No information is available on what the horses were fed pre-euthanasia, but it is conceivable that like the forages in this study, *Proteobacteria* was present in significant numbers. Epiphytic bacteria on forage may therefore have an impact on foregut bacterial profiles, but simultaneous profiling of feed and gut bacteria would have to be undertaken to determine the existence and strength of this relationship. Dougal *et al.,*[22,23] reported the presence of *Proteobacteria* in the equid gut but these were less abundant than *Bacteroidetes and Firmicutes* which were the major phyla found in the gut. S*iprochaetes, Actinobacteria* and *Fibrobacteres* were also present but to a lesser degree than the other 3 phyla. Thus, the four phyla that were represented by more than 90% of the bacteria in the hays in the current study i.e., *Proteobacteria, Cyanobacteria, Actinobacteria* and *Bacteriodetes* were present in the equid gut but at different proportions to that found in the gut. The fact that the equid core gut community, particularly that in the upper gastro-intestinal tract, which is composed of many small OTUs, lacks commonality between horses on similar diets, suggests that horse response to diet is unique and this could explain the susceptibility of some animals to digestive upset when fed similar diets to those that have no problems.

Clearly local environmental conditions contrive to favour the proliferation of some bacteria over others. For example, some bacteria such as *Enterobacteriaceae Rhodobacteraceae Bacillaceae Streptococcaceae*, while present in small numbers on dry leaves rapidly proliferated when wet, thereby altering the microbial profile of the leaf. Therefore, the distribution of common opportunistic bacteria together with the more specific residents to that particular phyllosphere under different environmental conditions can produce an ever-changing profile [52,53]. In hays this could be further altered by a pre-feeding treatment such as soaking.

The relationship between feed and foregut bacterial profiles in particular requires further investigation. The highly individualistic nature of the gut microbiome noted by both 25. Ericsson *et al.* [25] and Dougal *et al.,*[22,23], and the multiple small OTU core suggests that the gastric profile may lack robustness and be easily influenced by external factors. Understanding how the gastric microbiome responds to the epiphytic challenge from fodder may provide additional insight into gastric pathologies such as ulceration.

## Conclusion

The hays tested here supported a diverse population of epiphytic bacteria that was altered by pre-feeding soaking treatments. Grass type influenced the bacterial profile with multi-species meadow hays housing different profiles to the single species perennial ryegrass hay. The response to soaking was highly individualistic in terms of WSC leaching, bacterial numbers and profiles and there was no relationship between any of these parameters. While soaking for one particular time might be most effective for WSC reduction and produce little increase in bacteria in one particular hay, the results from this study show that a definitive recommendation on soaking duration cannot yet be made as other hay types could respond differently to the same treatment. Some increases in potential pathogens occurred post soaking and so caution should be employed when soaking fodder for stabled horses, particularly those with a previous history of colic. As hay is a major constituent of the horse’s diet and colic the major cause of death of horses across the developed world, additional studies on plant microbial communities, how pre-feeding treatments alter these and their interaction with the microbiome of the equid gastro intestinal tract are warranted.

## Acknowledgements

Thanks go to Leo Zeef University of Manchester for bioinformatics assistance and Sally Rice and Darren Hawkins for technical help, Royal Agricultural University.

